# Sandy: A user-friendly and versatile NGS simulator to facilitate sequencing assay design and optimization

**DOI:** 10.1101/2023.08.25.554791

**Authors:** Thiago L. A. Miller, Helena B. Conceição, Rafael L. Mercuri, Felipe R. C. Santos, Rodrigo Barreiro, José Leonel Buzzo, Fernanda O. Rego, Gabriela Guardia, Pedro A. F. Galante

## Abstract

Next-generation sequencing (NGS) is currently the gold standard technique for large-scale genome and transcriptome studies. However, the downstream processing of NGS data is a critical bottleneck that requires difficult decisions regarding data analysis methods and parameters. Simulated or synthetic NGS datasets are practical and cost-effective alternatives for overcoming these difficulties. Simulated NGS datasets have known true values and provide a standardized scenario for driving the development of data analysis methodologies and tuning cut-off values. Although tools for simulating NGS data are available, they have limitations in terms of their overall usability and documentation. Here, we present Sandy, an open-source simulator that generates synthetic reads that mimic DNA or RNA next-generation sequencing on the Illumina, Oxford Nanopore, and Pacific Bioscience platforms. Sandy is designed to be user-friendly, computationally efficient, and capable of simulating data resembling a wide range of features of real NGS assays, including sequencing quality, genomic variations, and gene expression profiles per tissue. To demonstrate Sandy’s versatility, we used it to address two critical questions in designing an NGS assay: (i) How many reads should be sequenced to ensure unbiased analysis of gene expression in an RNA sequencing run? (ii) What is the lowest genome coverage required to identify most (90%) of the single nucleotide variants and structural variations in whole-genome sequencing? In summary, Sandy is an ideal tool for assessing and validating pipelines for processing, optimizing results, and defining the costs of NGS assays. Sandy runs on Linux, MacOS, and Microsoft Windows and can provide feasible results, even on personal computers. Availability: Sandy is freely available at https://galantelab.github.io/sandy.

## 1. INTRODUCTION

The rapid development and cost reduction of next-generation sequencing (NGS) technology have made it the gold standard technique for large-scale investigations in genomics and transcriptomics (Shendure and Ji 2008; Levy and Myers 2016). The primary bottleneck in NGS is no longer the generation of sequences themselves, but the selection of appropriate computational tools and parameters to effectively analyze the generated data and extract accurate results from it (Zhao et al. 2017). Systematic evaluation of various approaches is a key strategy for addressing these difficulties. However, the task relies on having gold-standard NGS data with known “truth” values, which remains a challenge (Escalona et al. 2016).

Despite initiatives to generate well-characterized NGS data (MAQC Consortium et al. 2006; Zook et al. 2014), complete sets of true positives for experimental NGS data are unknown in practice (Escalona et al. 2016). Simulated or synthetic NGS (S-NGS) data have emerged as suitable solutions because they create a standardized scenario with well-defined true-positive values (Stephens et al. 2016). S-NGS data can benchmark the performance of different analytical methods for processing NGS data, tuning software, or pipeline parameters to obtain more accurate results or assist in experimental design (e.g., by leading the ideal sequencing coverage required for data analysis). However, overly simplistic simulated data that lack biological variations and common technical errors may be misleading and may indicate an underestimated amount of the data (e.g., sequencing coverage and number of replicates) required to achieve the expected results in an NGS assay.

Currently, many computational tools are available to simulate NGS data (Escalona et al. 2016; Zhao et al. 2017; Alosaimi et al. 2020). Curiously, most of these tools have been designed to simulate either genomic (Huang et al. 2012) or transcriptomic data (Griebel et al. 2012), but not both. More recently published algorithms that simulate whole-genome sequencing (WGS) data address the efficiency problem, add more complexity to simulations, and cover more sequencing platforms in response to the increasing complexity of genomic analysis (Henriksen et al. 2023; Yu et al. 2020; Nell 2020). Furthermore, highly specialized tools for simulating a specific type of data or sequencing platform are available (Li et al. 2018; Ono et al. 2013; Stöcker et al. 2016; Balzer et al. 2010). Moreover, recent reviews (Escalona et al. 2016; Zhao et al. 2017; Alosaimi et al. 2020) have pointed out that these tools have several limitations, including being difficult to use and computationally intensive, and lacking complete documentation, systematic benchmarks, or validations.

Herein, we present Sandy, a user-friendly and computationally efficient tool with complete computational methods for simulating NGS data from three platforms: Illumina, Oxford Nanopore, and Pacific Bioscience. Sandy requires only a reference FASTA file as input to mimic real NGS single/paired-end DNA or RNA sequencing. With parameters to control sequencing coverage, read length, sequencing error rate, and simulate reads with nucleotide variations or a real gene expression profile from several tissues, Sandy is highly flexible and adaptable to different scenarios.

## 2. RESULTS

### 2.1. Sandy’s pipeline: an easy to install, easy to use, and all-in-one tool for in-silico NGS assay

Sandy was designed following three main principles: i) to be easy to install and run in the most commonly used systems (Figure 1A), ii) to be of straightforward use and to simulate genomic and transcriptomic NGS data as similarly as possible to real assays (Figure 1B), and iii) to provide a complete set of parametrized steps to empower the user to mimic real genomic and transcriptomic data as best as possible (Figure 1C). Figure 1 outlines the main features of Sandy, including information about the simulation (e.g., for genome or gene expression), supported sequencing platforms, and sequencing type (Figure 1A and B). Figure 1C presents a more detailed version of the track that users can follow to simulate data, the dataset pre-included for genomics simulations (e.g., copy number variations (CNVs), single nucleotide variations (SNVs), and insertions/deletions (indels)), and transcriptome simulations (pre-computed gene expression data of 10k human samples of 52 tissues and 2 cell lines from genotype expression), which facilitate complex benchmark assays using Sandy.

**Figure 1.**
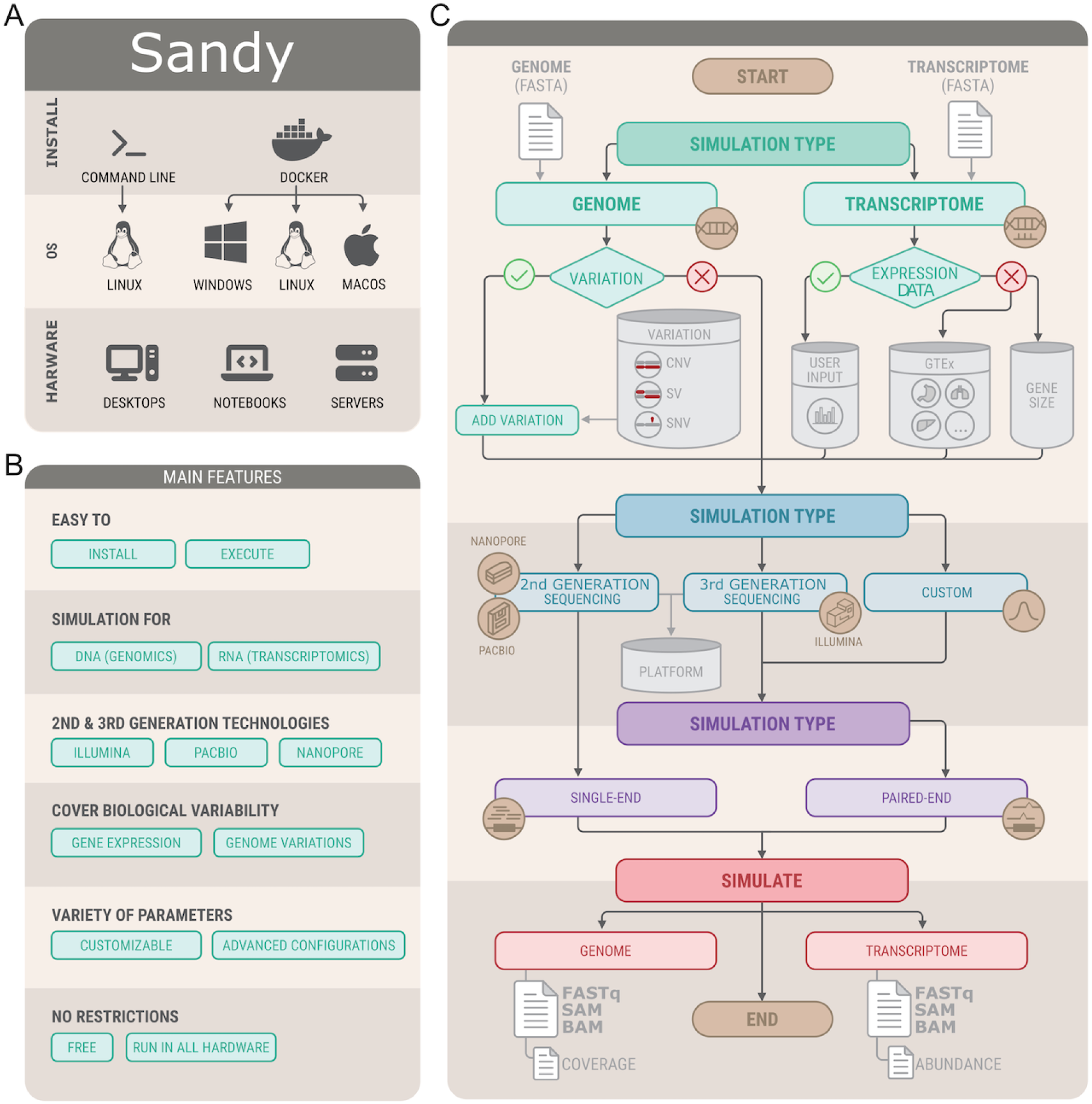
Sandy runs on all computer operating systems, is easy to use and install, and accurately replicates variabilities found in real NGS assays. A) Sandy can be directly installed or used as a Docker image, and it runs on the three most widely used computer operating systems with standard hardware such as personal computers and servers. B) An outline of all the main features available in Sandy. C) A schematic representation of Sandy’s pipeline, including simulation options, pre-computed and included datasets, and the primary resources available in Sandy.

To provide more details about each main Sandy feature, Figure 2 illustrates its practical use. Sandy is easy to install on the three most commonly used operating systems: Linux, Apple MacOS, and Microsoft Windows. Linux users can also directly install Sandy through the *cpanm* command-line (Figure 2). Sandy is easy to use because it runs on a command-line interface (CLI) and requires only a single input from the user: a genomic FASTA file for WGS simulation for Illumina, PacBio, or Oxford Nanopore, or a transcriptomic FASTA file for RNA sequencing (RNA-seq) simulations. For example, in Figure 2 (“Genome” box), Sandy simulates WGS on an Illumina HiSeq platform (paired-end reads of 101 bp). Figure 2 (“Transcriptome” box) it simulates RNA-seq run for 30 million paired-end reads of 101nt in length on the Illumina HiSeq platform resembling expression in the human liver.

**Figure 2.**
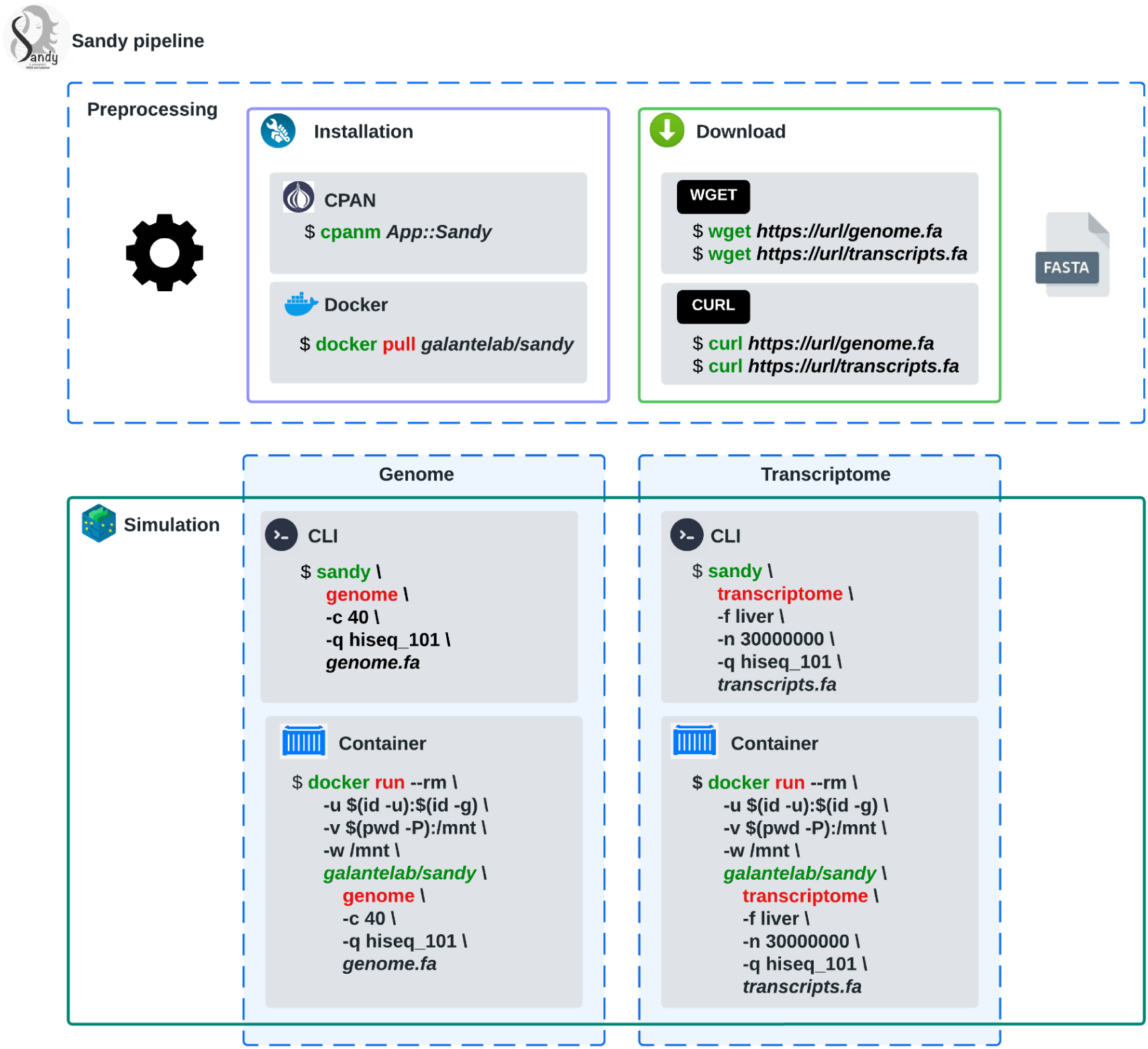
Installation and main commands on Sandy. Installation through Docker or CPAN. Example of usage for whole genome and transcriptome simulations with Sandy installed locally or using a Docker image.

Online Sandy documentation (https://galantelab.github.io/sandy/) provides a detailed explanation of its installation, uses it to simulate genomic and transcriptomic data, and provides numerous examples with comprehensive documentation.

### 2.2. Sandy simulation emulates variabilities found in real NGS assays

Sandy was designed to simulate NGS data as realistically as possible. It incorporates parameters and internal databases that consider common features and variabilities observed in NGS experiments. One of the key strengths of Sandy is its ability to simulate reads containing genomic variations commonly encountered in real NGS data, such as SNVs, indels, structural variations, and sequencing errors. For the RNA-seq simulation, Sandy included an internal database with gene expression profiles (52 human tissues and two cell lines) from Genotype-Tissue Expression (GTEx) data (GTEx Consortium 2013). This feature allows users to simulate the expression profile of a real human tissue (parameter-t). Moreover, users can upload their own custom expression profiles for use in the RNA-seq simulations. Detailed instructions can be found in the online documentation. Thus, simulations using different tissues (e.g., Liver and Testis) will result in reads from different sets of transcripts.

Sandy also mimics sequencing quality and sequencing errors, which can be selected from two options: empirical data (already integrated into Sandy or uploaded by users) or a Poisson model (see Methods, Discussion, and online documentation for details). Other platform-specific characteristics, such as fragment size (in paired-end Illumina sequencing) and variable read length (in PacBio and Nanopore sequencing), were also modeled and adjusted using Sandy’s parameters (Figure 1C). Additional parameters and descriptions are available in Sandy’s online documentation (https://galantelab.github.io/sandy/).

To demonstrate the similarities between the NGS data simulated by Sandy and the experimental NGS data, we conducted a comprehensive analysis by comparing the three features commonly analyzed in NGS assays. First, we examined the fragment length by WGS using paired-end reads on an Illumina platform (HiSeq). Figure 3A shows that Sandy’s data closely mimicked the experimental data in terms of fragment length. Next, we evaluated the base quality of the NGS reads. Figure 3B demonstrates that Sandy’s reads closely resembled the sequencing quality profile commonly observed in NGS data. Finally, we simulated RNA-seq datasets considering gene expression profiles in the Stomach, Testis, Pancreas, and Hippocampus and used these reads to quantify gene expression (Supplementary Figure 1). Figure 3C presents the principal component analysis (PCA) of gene expression between the simulated and experimental data (GTEx data). As expected, the gene expression patterns generated by Sandy closely grouped with their respective real tissues, confirming the accuracy of the expression simulation for each tissue type. In summary, Sandy effectively replicated the real characteristics and variability observed in experimental NGS data.

**Figure 3.**
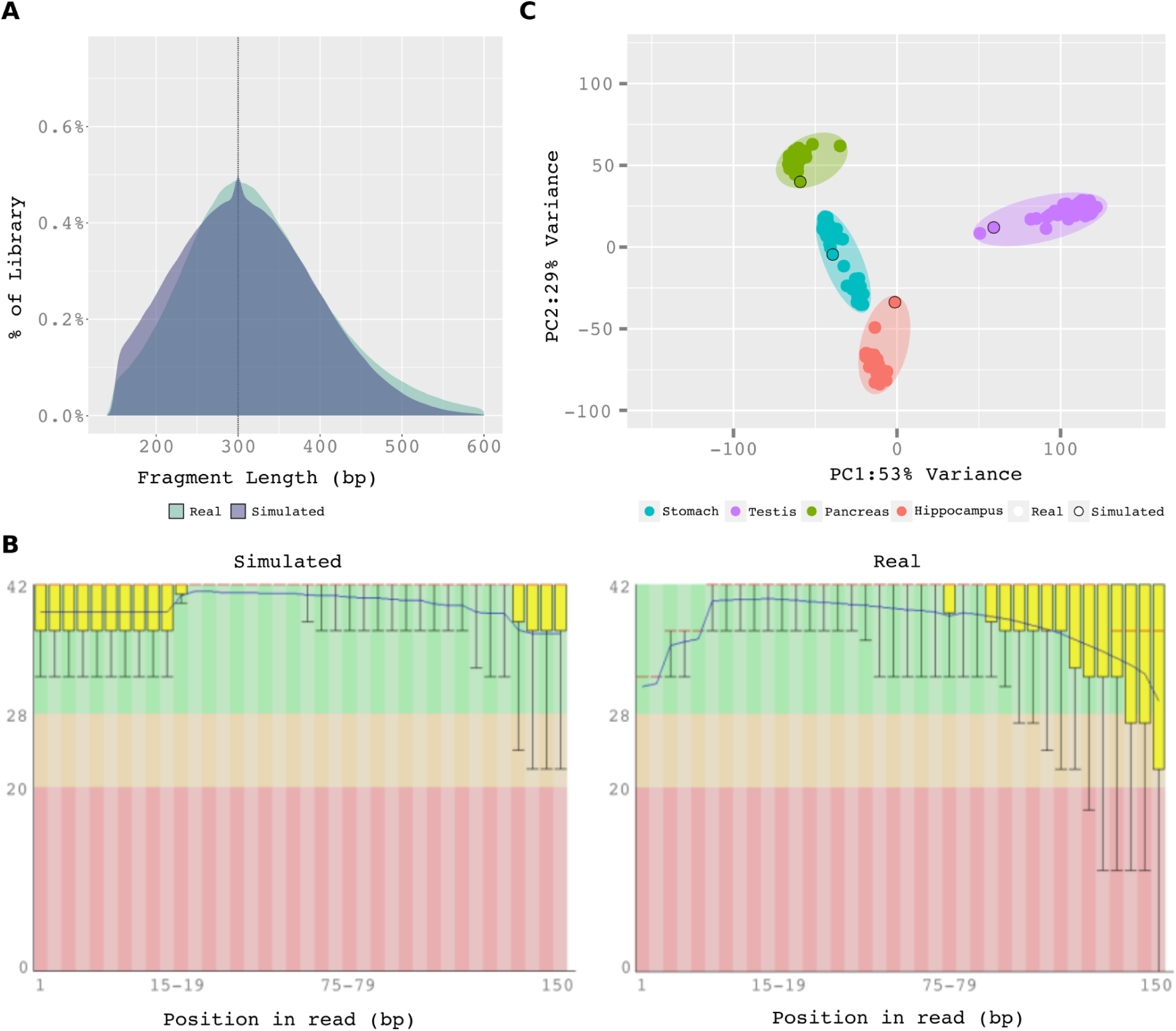
Simulated reads in Sandy closely resemble experimental NGS data. A) Distribution of fragment length in a real and simulated whole genome sequencing. B) FASTQ sequencing quality evaluation (real and simulated data are from an Illumina HiSeq platform). C) Principal component analysis (PCA) of gene expression from real (GTEx samples) and simulated (based on pre-computed gene expression included in Sandy) data from Stomach, Testis, Pancreas, and Hippocampus.

### 2.3. Output of simulations and data reproducibility using Sandy

Sandy’s output data are in the same format as those of the real NGS assay: FASTq (gzipped; -O gz; default), unaligned BAM (parameter: -O bam), or unaligned SAM (parameter: -O sam) (Figure 4A). Sandy generates files containing the expected values (positive control) associated with the simulation. The positive control for RNA-seq simulation is the expression count per transcript (Figure 4B, left side). For WGS, the number of simulated reads from each chromosome is used as the positive control (Figure 4B, right side). Another useful resource in Sandy that could be used as a positive control is the simulated read name (ID). It contains the transcript ID from which the read was simulated in an RNA-seq simulation (Figure 4C, left side) or the genomic chromosome, orientation (from leading (P) DNA strand or lagging (M) DNA strand), and start-end position in the genome for genome sequencing simulation (Figure 4C, right side).

**Figure 4.**
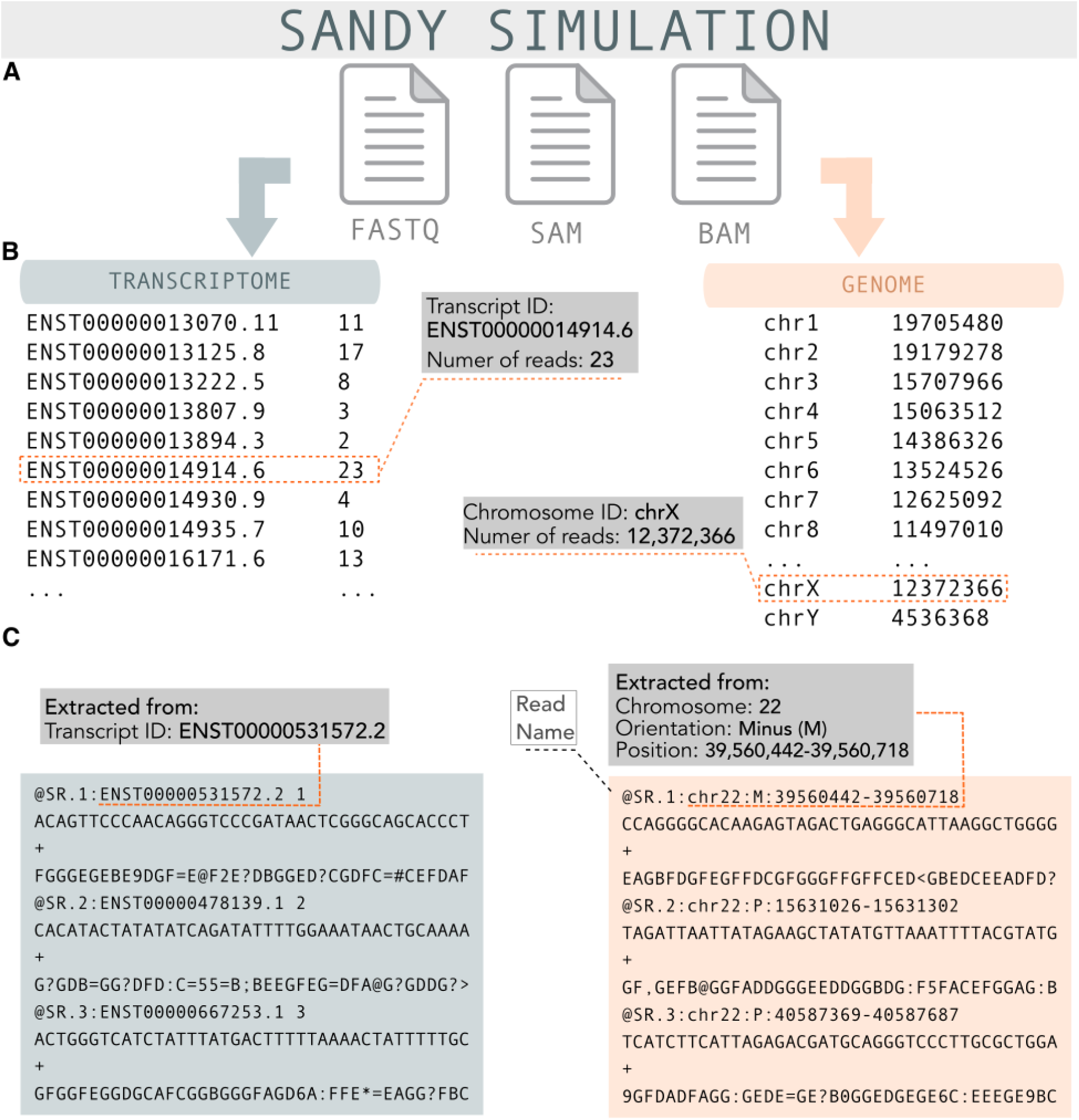
Sandy output includes FASTq, unaligned BAM, or SAM and practical positive control files. A) Three output formats are available for genome or transcriptome simulations performed in Sandy: FASTq, SAM, and BAM. B) Expression counts per transcript are provided for simulations of RNA-seq, while the number of simulated reads from each chromosome are provided for simulations of whole-genome sequencing. C) Transcript IDs from which reads were simulated are provided for RNA-seq simulations, while genomic position and orientation are provided for whole-genome simulations.

Reproducibility is a major issue in computational biology (Papin et al. 2020). Therefore, Sandy has the option “*--seed*” that receives an integer used to initiate the random number generator. Additionally, users must set the same number of jobs as stated in the previous simulation, as each job receives a different seed calculated from the main seed and, obviously, all other chosen parameters.

### 2.4. Performance: Sandy runs in all types of hardware

In practice, methods that require powerful computational structures are either prohibitive or limited for most researchers. To avoid such limitations, Sandy was optimized to use hardware efficiently (e.g., using multiple threads) while consuming minimal memory and CPU time. Sandy runs well on standard laptops or desktop (e.g., Intel i5/i7 cores; 8–16 GB RAM; 0.5–1 TB storage) or standard servers (e.g., Intel Xeon with 12 cores; 64 GB RAM; 1 TB storage). To demonstrate Sandy’s performance, we executed various RNA-seq and whole-genome simulations using a *laptop* and *server* (details provided in Methods and Supplementary Figure 2-3). Figure 5A shows the runtime for RNA-seq simulations on the Illumina HiSeq platform with varying numbers of reads, ranging from 10 to 150 million paired-end reads (Supplementary Figure 2). Even at the highest end of the simulation (150M), Sandy completed the task within a reasonable timeframe of approximately 1 h (67 min). On the laptop, the runtime was still feasible, with a simulation of 50 million reads in less than 2 h (1 h and 44 min).

**Figure 5.**
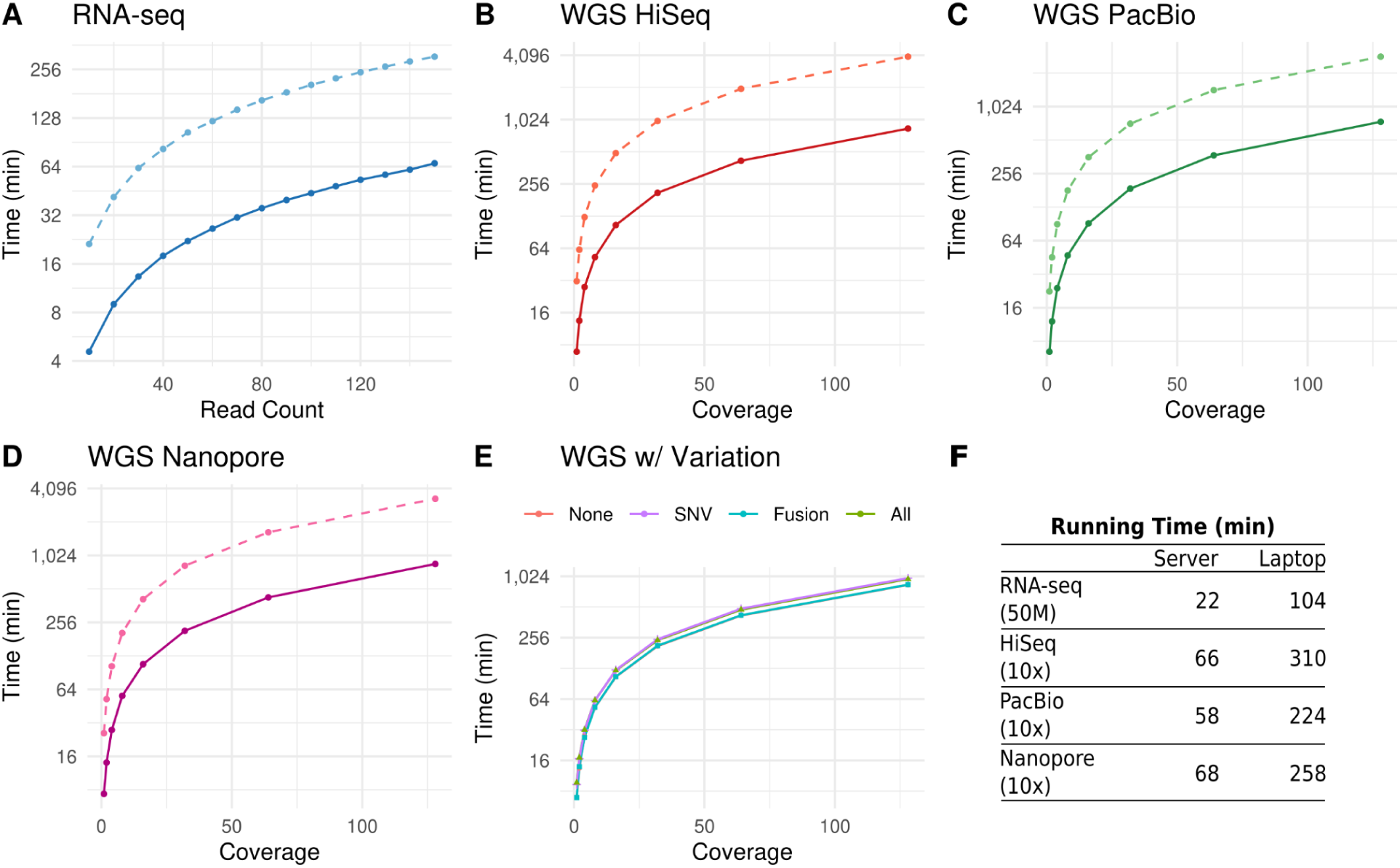
Sandy can efficiently operate on standard servers and laptops. Simulation time of RNA and whole-genome sequencing (time in log2 scale). Dashed lines represent execution times obtained from the laptop, whereas full lines refer to execution times on the server. (A) Time variation dependent on read count simulation (RNA-seq). (B) Time variation dependent on simulated coverage for Illumina HiSeq. (C)Time variation dependent on simulated coverage for PacBio. (D)Time variation dependent on simulation coverage for Oxford-Nanopore. (E) Time variation dependent on simulated structure variation for Illumina HiSeq. (F) Execution times of common simulations in genomics and transcriptomics.

WGS simulations were performed with coverage varying from 1X to 128X based on the Illumina (HiSeq), Nanopore, and PacBio platforms for the human genome (Supplementary Figure 3). Figures 5B–D show Sandy’s performance on both the server and laptop. A WGS simulation with approximately 30X coverage (approximately 90 billion bases, on average, producing 90 GB of data) required less than 4 h (3 h and 31 min) (Illumina HiSeq). Even for an extremely high coverage of 128X (Illumina HiSeq), the simulation required less than 15 h (14 h and 3 min). As expected, when running the simulation on a laptop, the execution time was, on average, 4.5 times longer than that on the server. However, this is not prohibitive and only reflects the hardware limitations that users may encounter (Figure 5B–D). We also quantified Sandy’s performance by considering the inclusion of SNVs, gene fusions, or both. Remarkably, we did not observe an increase in the running time (∼1.17 fold on average) when these additional parameters were included (see Methods, Section 4.1.3 for more details). In summary, Figure 5F indicates how well Sandy performs (time) for some of the most common simulations in genomics and transcriptomics (RNA-seq: 50M reads; WGS: human genome 10X coverage).

### 2.5. Using Sandy to optimize read coverage in an RNA-seq assay

Sandy is a tool with many practical applications, including addressing a common and cost-conscious question in RNA-seq assays: What is the minimal number of reads required to achieve unbiased gene expression quantification for my samples? To answer this question, we conducted RNA-seq simulations with varying numbers of read pairs (10M, 30M, 50M, 100M, and 150M) for different tissues using the expression profile (from GTEx) stored in the Sandy database for the Brain Cortex, Hippocampus, Liver, Pancreas, Bladder, and Stomach (Supplementary Figure 4 shows all the command lines used for this simulation). Figure 6A–C shows the correlation plots between the expected counts of protein-coding transcripts expressed in pancreatic tissue, as in GTEx samples, and the observed counts simulated by Sandy and quantified by Kallisto (Bray et al. 2016). A positive correlation was observed between the real and simulated data (R²). Figure 6D summarizes how R² behaves in different tissue types depending on the number of reads in the simulated RNA-seq experiment. Interestingly, we observed that increasing the number of reads up to 50M reads rapidly increased the accuracy quantification (from an average of R² ∼ 0.75 to R² ∼ 0.87) and that the gain was more modest for simulations with a higher number of reads, where we see that intrinsic biological factors possibly start to affect the accuracy. We also used simulations to estimate the minimum sequencing depth required to better estimate the lncRNA expression (based on lncRNAs annotated in Gencode v36; see Methods for details and Supplementary Figure 4). Notably, the quantification accuracy for lncRNAs exhibited a more rapid improvement compared to that of protein-coding transcripts as the sequencing depth increased. This improvement remained substantial for sequencing depths higher than 50M reads, making it a crucial consideration for experimental design. Not only is the number of lncRNA transcripts smaller in all tissues analyzed (approximately 7,000 transcripts with expression >0.1TPM), but the expression of these transcripts is also lower by one order of magnitude than that of protein-coding transcripts, making them highly sensitive to an increase in sequencing depth. Notably, Sandy used gene expression data from the GTEx consortium, which may have limited the coverage of non-polyadenylated lncRNAs. Altogether, Sandy simulations can also lead researchers to refine their experimental designs in NGS assays, improve downstream data analyses, and optimize their budgets.

**Figure 6.**
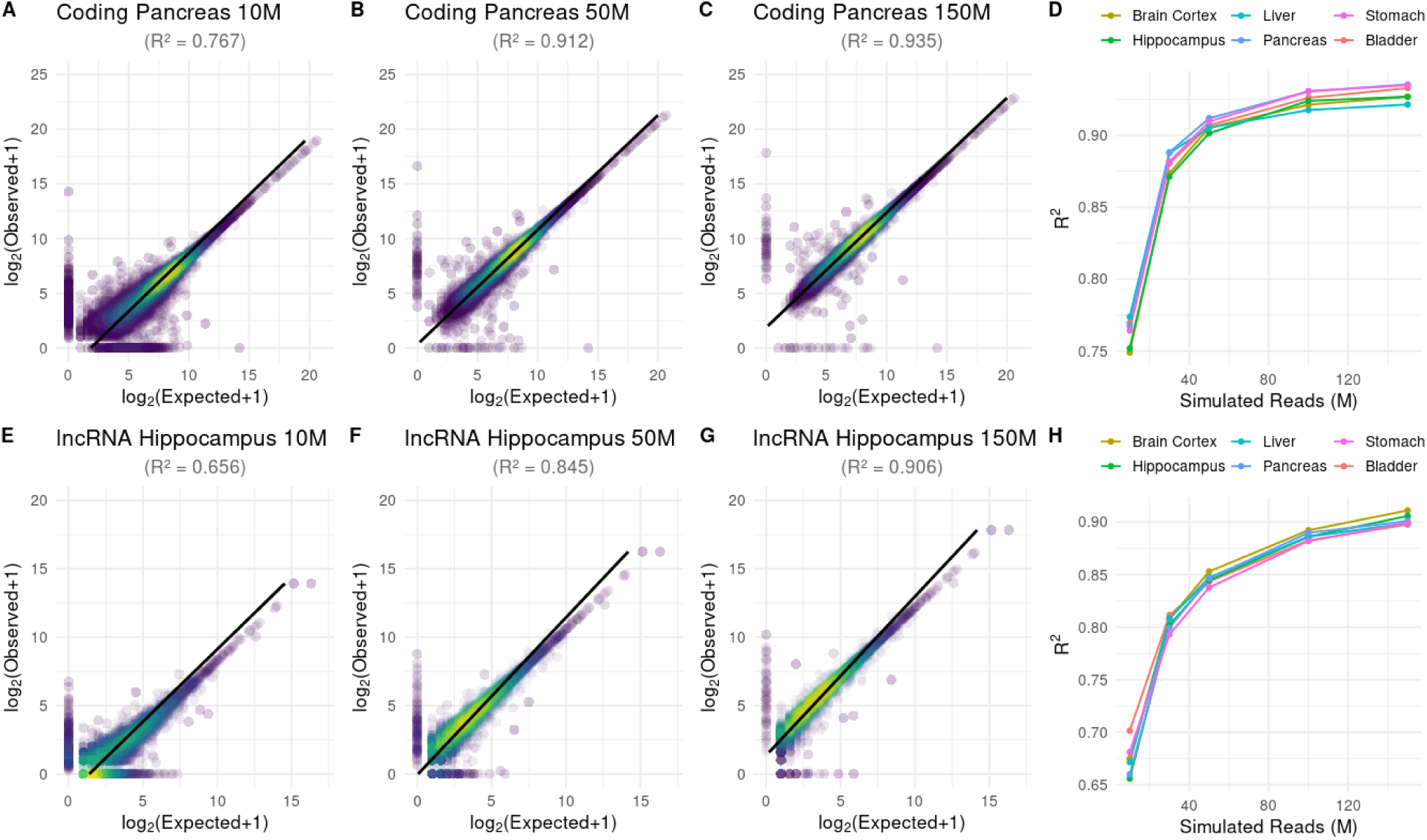
Sandy can be used to estimate the required number of sequencing reads for an RNA-sequencing assay. A–C and E–G show correlation plots of expected (in real data) reads and observed reads (quantified after simulation). (A) 10 million simulated reads of coding RNA for Pancreas; (B) 50 million simulated reads of coding RNA for Pancreas; (C) 150 million simulated reads of coding RNA for Pancreas; (D) R² variation dependent on the number of reads (in millions) for coding RNA according to different tissues; (E) 10 million simulated reads of lncRNA for Hippocampus; (F) 50 million simulated reads of lncRNA for Hippocampus; (G) 150 million simulated reads of lncRNA for Hippocampus; (H) R² variation dependent on the number of reads (in millions) for lncRNA according to different tissues. All quantifications were performed using Kallisto and Gencode v36.

### 2.6. Using Sandy to optimize the genome coverage in an NGS assay

Subsequently, Sandy was used to estimate the optimal cost-benefit of genome sequencing coverage for detecting SNVs, indels, and structural genomic variations in the WGS dataset (Supplementary Figure 5). Specifically, we used Sandy to simulate WGS coverage of 4X to 128X (for simplicity, we analyzed only chromosome 1). Additionally, we used Sandy’s parameter-*-genomic-variation* to include SNVs and indels (Supplementary Figure 5) in individual NA12878. NA12878 SNVs (http://www.internationalgenome.org/) for all chromosomes are currently available for Sandy and can be easily called using a simple parameter option. Next, we used the GATK toolkit https://gatk.broadinstitute.org/ for variant calling (and methods) and compared the results with the expected SNVs. Briefly, variants that were represented in the simulated reads and called by GATK were considered true positives, variants absent in the simulation but called by GATK were considered false positives, and variants in the simulated reads but not called by GATK were considered false negatives. As expected, genome coverage and variant detection were positively correlated (Figure 7A). In the low-coverage simulation (4X and 8X), less than 70% of the true-positive SNVs were called. However, with 30X coverage, more than 90% of SNVs were accurately identified (Figure 7A). A plateau of calling was achieved at approximately 60X and little gain in terms of SNVs was obtained over this limit. Interestingly, the false positive calling was from specific genomic regions, especially regions near centromeres/telomeres (Figure 7B) and regions with notably poor mapping ability.

**Figure 7.**
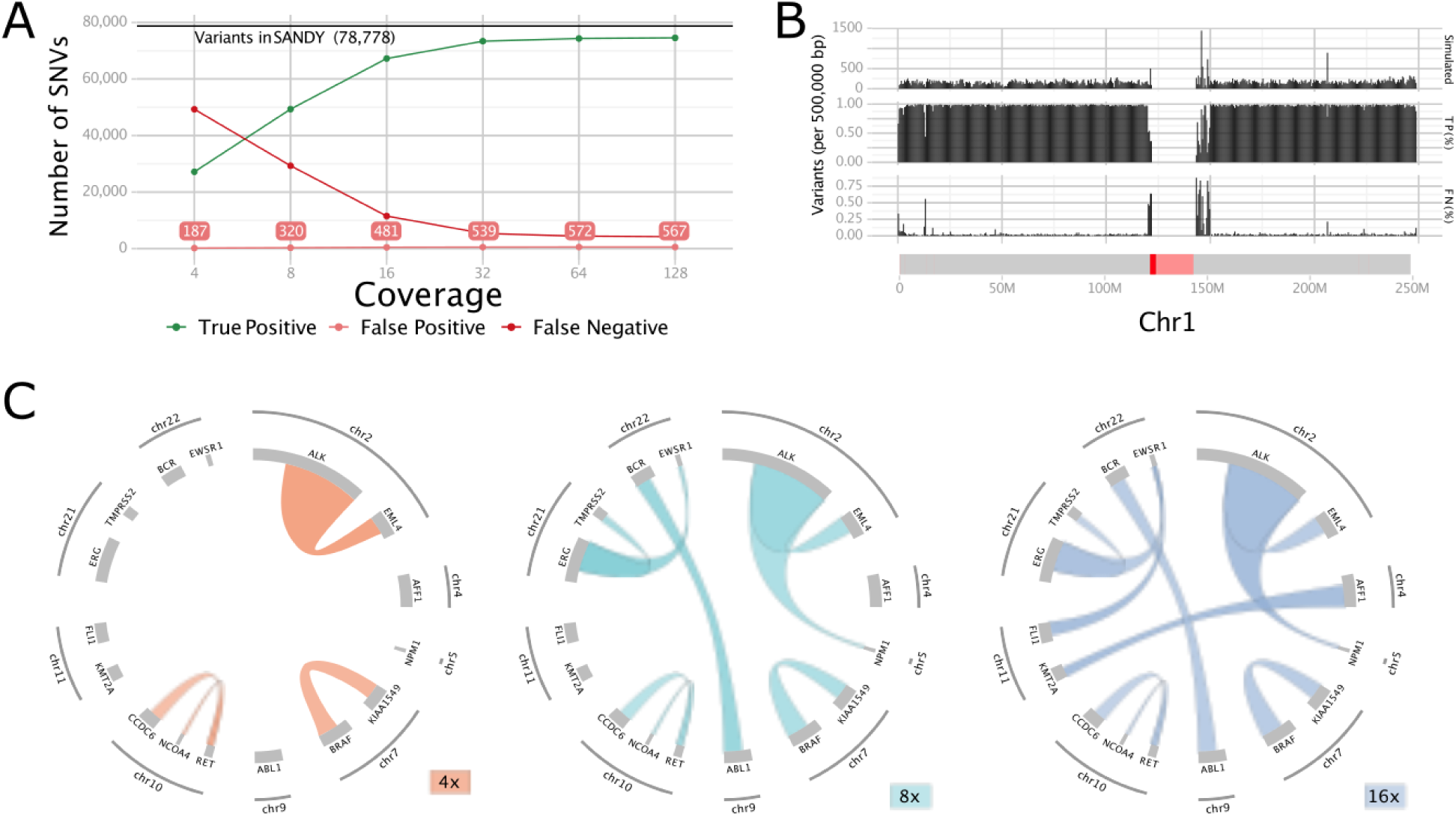
Sandy can be used to estimate the optimal coverage for accurate genomic variation identification. A) Variant calling performance for simulations with varying sequencing coverage. The y-axis shows the number of variants simulated and found (true positive), number of variants simulated and not found (false negative), and number of variants not simulated but found (false positive). B) Distribution of single nucleotide variations (SNVs) simulated in chromosome 1. Top: distribution of simulated variants; middle: percentage of true positives; bottom: percentage of false negatives. The X-axis represents chromosome 1, where the centromere annotation is in dark red and not sequenced centromere regions (identified as “Ns” in the reference genome) are in light red. C) Circos plot showing gene fusions detected at coverages of 4X (40%), 8X (70%), and 16X (all).

Regarding structural variations, we performed WGS simulations at coverages of 4X, 8X, and 16X, including ten cases of homozygous gene fusion (Supplementary Table 1; Supplementary Figure 6). These fusions are common cancer gene fusions, such as BCR-ABL1 (Philadelphia chromosome, mainly found in chronic myeloid leukemia), EML4-ALK (mainly found in lung cancer), CCDC6-RET (found in thyroid papillary carcinoma), and EWSR1-ERG (Ewing sarcoma), and are native to Sandy (parameter *-a*). As expected, the detection of gene fusions positively correlated with sequencing coverage. Specifically, at 4X coverage, 40% of the gene fusions were identified; at 8X coverage, 70% were detected; and at 16X coverage, all gene fusions were correctly identified (Figure 7C and Supplementary Table 1). It is important to note that we did not include tumor heterogeneity in this simulation. The versatility and effectiveness of Sandy lie in its capacity to seamlessly simulate NGS experiments that account for various genomic structural variations and SNVs in a single command line. Sandy output is readily processed by state-of-the-art algorithms used routinely in NGS data analysis and yields detection rates as expected from real data, reinforcing our belief that Sandy will be an invaluable tool for researchers when testing and fine-tuning existing bioinformatics tools as well as in the development of novel pipelines for detecting genomic variation.

## 3. DISCUSSION

Here, we present Sandy, a tool that simulates genomic and transcriptomic data for the Illumina, PacBio, and Nanopore platforms. Sandy was designed not only to be easy-to-install and easy-to-use, but also as a flexible tool capable of running in standard hardware and on the most common operating systems, being complete in terms of resembling a real NGS assay. NGS is the leading technology for exploring genomes and transcriptomes and is used to answer increasingly demanding questions in biology and medicine (Goodwin et al. 2016). Many software tools have been developed to process NGS data, each with its own distinct errors, biases, and low concordance (Kumaran et al. 2019; Barbitoff et al. 2022). Thus, careful benchmarking is required to assess the performance of existing and newly designed computational genomic pipelines and to quantify their sensitivity and specificity for any given study. However, important bottlenecks in the pre-experimental design, especially in post-data processing, are common in NGS assays (Zhao et al. 2017). The use of NGS with known “true” values (positive controls), which can be used for benchmarking pipelines of data analyses, is a solution to overcome these bottlenecks. In contrast to experimental data, in which the true values are often unknown, simulated datasets are generated with predefined attributes in a standardized context. Although some empirical datasets are available for this purpose (MAQC Consortium et al. 2006; Zook et al. 2014), they are specific to certain assays and not available for many methods and organisms. Therefore, simulators of synthetic sequence data, such as Sandy, offer a controlled environment in which the “ground truth” is known. These simulators may be essential tools for better defining NGS assays and their processing pipelines.

To date, many NGS simulators have been developed (Escalona et al. 2016; Alosaimi et al. 2020; Zhao et al. 2017). However, as recently noted by Escalona et al., these tools have some limitations. For example, i) they have not been extensively benchmarked or validated. Approximately half of these tools have been described in sections of another article without a dedicated publication; ii) the pipelines are largely redundant, implementing the same or very similar functionalities; iii) they are poorly documented and can be difficult to use by non-experts; and iv) some of them are no longer maintained. In contrast, Sandy comprehensively overcomes these limitations. This article provides a full and detailed description of our tool, ensuring transparency and providing a comprehensive understanding of its functionalities. Extensive online documentation is available (https://galantelab.github.io/sandy) to show how Sandy can be used by any researcher, regardless of expertise. Our tool combines novel and useful functionalities, such as the gene expression profile for RNA-seq simulation, simulation of genomic variations, base quality estimation based on real data, and simulation of second-and third-generation NGS platforms. Sandy is easy to use and requires only a single FASTA file as the input in a streamlined command line. It can be installed easily (Docker-encapsulated image or command line) and is fully available under a GNU license on GitHub. Since its creation, we have improved Sandy seven times (current version: 0.24) and will keep updating Sandy frequently (at least once a year).

To demonstrate the utility of Sandy, we addressed the critical considerations underlying NGS experiments: i) in a RNA-seq assay, to determine the optimal number of reads for the RNA-seq to accurately quantify gene expression profiles; ii) in WGS, to establish the appropriate genome coverage for capturing the genomic variations of interest. Researchers often estimate these values based on previous experience or values reported in other studies. For instance, it is widely accepted that an RNA-seq experiment requires a minimum range of 30–50 million reads to effectively quantify gene expression (Conesa et al. 2016), and a 30X fold coverage is typically used in WGS to identify genomic variations. However, these general guidelines may overlook the inherent complexity of individual samples. For example, certain tissues, such as the brain, have a higher number of expressed genes and isoforms than the human liver, demanding deeper sequencing. Similarly, sequencing depth may vary based on the specific set of genes of interest (protein-coding vs. long non-coding). In this context, we demonstrated how easily Sandy can simulate NGS data that emulate biological complexity, such as gene expression patterns. By utilizing Sandy, researchers can better design their NGS experiments and tailor the sequencing depth and coverage to suit the specific complexities of their samples. Moreover, Sandy simulation capabilities enable researchers to fine-tune their pipelines and parameters, optimize the analysis of NGS data, and enhance the accuracy of their findings.

As expected, Sandy has limitations that can be improved. First, Sandy did not simulate all the current sequencing platforms, such as the Ion Proton, Complete Genomics, and Helicos platforms. Second, context-specific methodologies such as single-cell RNA-seq, ATAC-seq, Hi-C, and ChIP-seq are not covered in Sandy simulations. Third, Sandy does not have a pre-computed database for other species, but only for humans. Fourth, Sandy’s options for mimicking the complexity of real biological data and their variability (e.g., the genomic heterogeneity found in cancer) are far from complete. We understand the importance of providing additional options for researchers to generate trustworthy synthetic data, and several of Sandy’s limitations can be overcome in future studies. In the current version of Sandy, users can upload pre-computed datasets (e.g., gene expression and sequencing quality); however, further pre-computed datasets for other species, pathological conditions (e.g., gene expression profiles from cancer), and heterogeneity in genomes will be included in future versions of Sandy.

More broadly, there is no NGS simulator capable of fully capturing all the experimental variabilities and noises (e.g., batch effects) found in an NGS assay. For example, in practice, a cancer sample contains not only tumor cells (which are heterogeneous per se) but also tumor-associated normal epithelial and stromal cells, immune cells, and vascular cells, which increases the set of genome variants and complexity in terms of gene expression. Thus, all NGS simulators, including Sandy, underestimate some features of the experimental NGS assay.

Despite these limitations, in silico simulation of NGS has emerged as a powerful tool to improve NGS analyses and may help improve the design of NGS assays and processing pipelines. We are convinced that Sandy is a valuable tool for researchers given its ease of installation, ease of use, versatility, and customization. Sandy functions seamlessly on standard hardware and common operating systems and is a more complete tool with functionalities that have never been combined before in a pipeline to simulate NGS data.

## 4. METHODS

### 4.1. Sandy: algorithm implementation

Sandy was implemented in Perl (https://www.perl.org/) and was structured into five main modules: i) simulator, ii) read, iii) quality, iv) database manager, and v) auxiliary components (Figure S1). These modules depend heavily on the freely available modules of the Comprehensive Perl Archive Network (CPAN, https://metacpan.org/). Sandy’s source code is freely available under the GNU General Public License at https://github.com/galantelab/Sandy/blob/master/LICENSE.

#### 4.1.1 Input

Sandy processes nucleotide sequences in the FASTA format as input. Sequences can be labeled as transcripts or chromosomes. When a user requests a simulation from Sandy, the auxiliary component *command line parser* is first called to process (parse) a user request. Subsequently, the *parameter checking* component verifies and validates all the requested parameters. Next, the *simulator* module is called. This module is responsible for genome and transcriptome sequencing simulations, and is structured into multiple components. The first component is the *index FASTA*, which validates the submitted FASTA file and stores the FASTA sequences in a hash table data structure. The *strand selector* component is then set to select a strand randomly at any time it is called. The DNA strand (plus or minus) is selected based on the type of the simulation. Genomic simulations randomly sample the strand with a 50% chance for each strand, whereas transcriptomic simulations are based on cDNA sequencing; therefore, only the minus strand is used.

#### 4.1.2. Simulating read distribution

Sandy was implemented by using a negative binomial distribution to model the number of reads generated by each sequence. The *sequence selector* component randomly selects the chromosomes/transcripts to achieve the required distribution. By default, the sequence lengths are used for weighted sampling of positions within the given FASTA sequences. Alternatively, for transcriptome simulations, Sandy allows users to input an expression matrix or select an expression matrix from Sandy’s internal database (for more details, see Section 4.1.6). The *expression checking* component recovers the requested expression matrix (if any) from the Sandy database and stores it in a hash table. This feature provides flexibility for users to create simulations with certain tissue types or with specific biological and technical models.

#### 4.1.3 Simulating genomic variations

Users can request the inclusion of genomic variations using Sandy’s native database. Sandy makes available SNVs and indels from individual NA12878 and fusions common in cancer genes (more details in Section 4.1.6). Users can add the desired variations in vcf file format to Sandy’s database using a simple command line (extensive details can be found on Sandy’s webpage). When a genome simulation is requested, the *variation checking* component recovers the requested variations (if any) from Sandy’s database and stores them in a piece table. To avoid retaining various FASTA versions in computer RAM, only the alterations (as insertion and deletion coordinates) are loaded into a read-only list of pieces, leaving the original sequence vector intact. Genomic alterations are applied immediately after sequence raffling, keeping the piece list instructions, following the final assembly, and generating a new simulated fragment/read.

#### 4.1.4 Simulating NGS protocols

Sandy can perform simulations on three main platforms: Illumina, Nanopore, and PacBio. For Illumina simulations (WGS or RNA-seq), users can use standard quality profiles for HiSeq, MiSeq, or NextSeq with default mean *read* length, standard deviation, and *error*. In addition, these parameters can be set by users to better replicate NGS assays. For Nanopore and PacBio, the most challenging issue is to define the read length because it may vary from less than a hundred to more than a thousand. In Sandy, the read lengths for Nanopore and PacBio were defined based on the distribution of values obtained from real experiments performed on these platforms. For PacBio, the read size window was defined between 5,000 and 12,000nt on average (Zook et al. 2014), whereas for Nanopore, the values ranged between 8,000 and 21,000nt on average (Jain et al. 2018).

The *read* module is responsible for assembling the single/paired-end reads. It receives the chromosome/transcript sequence and randomly samples the position from which it takes a sequence slice according to the fragment size. Fragment lengths are drawn from a normal distribution with mean size and standard deviation. By default, the fragment mean size is 300nt and the standard deviation is 50nt, but users can change these parameters. The sliced sequences then undergo some variations based on the sequencing error rate and genomic variations from the *variation* component. Reads are generated from the sliced sequences using the *read_single* (single-end reads) or *read_paired* (paired-end reads) component.

#### 4.1.5 Base quality

Sandy models base quality following two standalone models: i) a Poisson distribution, which works for all read lengths (Bliss and Fisher 1953); and ii) a quality-score distribution extracted from empirical data, which is available for some Illumina read lengths, all Nanopore, and all PacBio reads. Sandy mimics real sequencing experiments by preprocessing the Illumina (data from (Fairley et al. 2020)), PacBio (data from (Zook et al. 2014)), and Nanopore (data from (Jain et al. 2018)) datasets. Users can upload sequencing-quality data from their own platforms to better simulate the desired NGS assays. The *quality* module recovers the quality profile from Sandy’s database and stores it in a Phred score vector for each quality sequence. To generate a quality string, the module samples each score from the vectors of the Phred scores until the required read size is reached. For a theoretical Poisson-quality entry, a quality string is generated by raffling each position from a Poisson distribution.

#### 4.1.6 Databases

Sandy has three internal databases (*variation*, *expression*, and *quality*) implemented in SQLite3 (https://www.sqlite.org/index.html). They count on selected default entries as follows: i) base quality profiles for Illumina, PacBio, and Nanopore sequencers; ii) genomic variations from the NA12878 genome (SNVs and indels) and gene fusions from COSMIC (Tate et al. 2019); and iii) gene expression profiles from 54 GTEx tissues (GTEx Consortium 2013). In addition, the user has the option to add their own entries, thus customizing the genomic/transcriptomic sequencing simulations.

The *database manager* module is responsible for managing the user-defined extensions in Sandy’s database. When a user attempts to dump a new entry into Sandy’s database, the auxiliary component *command line parser* is called to process the user request first, and the *parameter checking* component verifies and validates the requested parameters. The dataset to be dumped is validated by the *validation* component according to each database, indexed by the *index extension* component, and added to the SQLite3 database by the *add to database* component.

#### 4.1.7 Writing output

Once Sandy generates sequencing reads, the *template* component is finally set to either SAM or FASTQ (or their compressed versions, i.e., BAM or gzipped FASTQ) based on user choice. The reads are written in user-specified format. Sandy also generated files containing the expected values associated with the simulation, that is, expression count per transcript for RNA-seq simulation and number of simulated reads per chromosome for WGS, facilitating a positive control in downstream analysis. For each simulated read, Sandy automatically generates a sequence identifier containing the sequence ID from which it was simulated. For the WGS simulation, the sequence identifier also contains the orientation and position of start ends in the genome (section 2.3). Users can also customize the sequence identifiers.

#### 4.1.8 Simulation reproducibility

Sandy has the “*--seed*” option, which receives an integer used to initiate the random number generator. In addition, users must set the same number of jobs because each job receives a different seed calculated from the main seed. Therefore, simulations with the same parameters, seeds, and number of jobs are identical.

#### 4.1.9 Runtime and data storage

Sandy was developed with a multicore threading framework; therefore, the runtime decreases as more cores are allocated. Time and hard disk storage scale linearly with the number of simulated reads, independent of the type of platform or type of simulation (RNA-seq or WGS) (Section 2.4). Because of the strategy adopted in Section 4.1.3, the execution time does not increase substantially when genomic variation is added, which is a remarkable feature of Sandy.

### 4.2. Performance evaluation and simulation of WGS and RNA-seq data

To evaluate Sandy’s performance, WGS and RNA-seq simulations were executed in both a standard laptop (Intel(R) Core(™) i5-8250U CPU @ 1.60 GHz, 8 GB RAM, 256 GB HD, running a Linux operating system) and a server (Intel(R) Xeon(R) CPU E5649 @ 2.53 GHz, 128 GB RAM, 1 TB HD, running a Linux operating system). Specifically, the wall clock time was determined using the Unix “time” command. Only basal computational processes were maintained during the performance assessments.

For all WGS performance evaluations, paired-end reads were simulated for the human genome (hg38/GRCh38) using Sandy’s Docker image. Simulations were performed for coverage varying from 1X to 128X, and according to three quality profiles: Illumina HiSeq (101 paired-end reads), Nanopore, and PacBio. In addition, we performed WGS simulations considering four scenarios: (i) no genomic variation, (ii) all NA12878 SNVs/indels, (iii) 10 gene fusions, and (iv) both SNVs/indels and gene fusions. All the genomic variations were directly included in Sandy’s native database. To evaluate RNA-seq performance, we used the expression profile of human liver tissue available in Sandy’s database. We simulated paired-end reads by varying the read depths from 10 to 150 million reads (for every 10 million reads). All the remaining parameters were considered to be default.

### 4.3. Gene expression analyses

To assess gene expression on simulated data, we first simulated RNA-seq reads based on human GENCODE v36 transcripts using Sandy’s Docker image for the Illumina HiSeq quality profile (76 bp paired-end reads), paired-end, fragment-mean = 237nt, and fragment standard deviation = 82nt. The datasets were simulated based on the expression profiles of the human Bladder, Hippocampus, Brain Cortex, Liver, Pancreas, Stomach and Testis available in Sandy’s database at read depths of 10, 30, 50, 100, and 150 million reads. The resulting FASTQ files and expression matrices, named *expected values*, were stored for further analysis. Next, the gene expression of FASTQ files for a coverage of 100M was assessed using the GTEx pipeline (https://github.com/broadinstitute/gtex-pipeline/blob/master/rnaseq/README.md) based on the human GENCODE v26 annotation. The FASTQ files for all coverage were quantified using Kallisto (version 0.48) based on the human GENCODE v36 annotation. Kallisto outputs were labeled as *observed values* and linear correlations with the *expected values* (obtained from Sandy’s expression matrices) were analyzed. Pearson R^2^ coefficients were obtained for the expected pairs of protein-coding transcripts (at least one pair with expression greater than 1TPM) and long non-coding transcripts (at least one pair with expression greater than 0.1TPM) and plotted as a function of sequencing depth for all simulated tissues.

### 4.4. Gene fusion detection

To identify the gene fusions reported in this manuscript, we first simulated WGS reads for the human genome (hg38/GRCh38) using Sandy’s Docker image. Simulations were performed according to the Illumina HiSeq quality profile (101 paired-end reads) with varying coverages (4X, 8X, and 16X), and included 10 gene fusions from Sandy’s native database (e.g., parameters -q hiseq_101 -j 20 -c 4 -v -i ’variation=%v’ -A fusion). For each simulated dataset, all reads were mapped against the human reference genome using BWA-MEM (Li and Durbin 2009) (parameters t 10 -T 0 -R “@RG\tID:GROUP1\tSM:SIM4x\tLB:SIM4x_LIB\tPL:ILLUMINA”). Subsequently, only reliable alignments against the genome (mapping quality score (Q) ≥ 20) were selected and sorted using SAMtools (Li et al. 2009). Duplicate reads were marked using Picard MarkDuplicates (http://broadinstitute.github.io/picard/) and high-quality reads were analyzed using Delly (Rausch et al. 2012) (version: 0.7.8; standard parameters) to identify potential structural variants (SVs), in particular the germline gene fusions. Finally, gene fusion candidates were filtered to remove low-quality predictions (paired-end support (PE) < 3 or median mapping quality of paired ends (MAPQ) < 20).

### 4.5. SNV/indels calling

We simulated the genome sequencing of CRC hg38’s chromosome 1 with varying coverages (4X, 8X, 16X, 32X, 64X, and 128X) using Sandy’s Docker image with a custom set of variants (added to Sandy as `chr1`). The simulation was performed using the following commands:

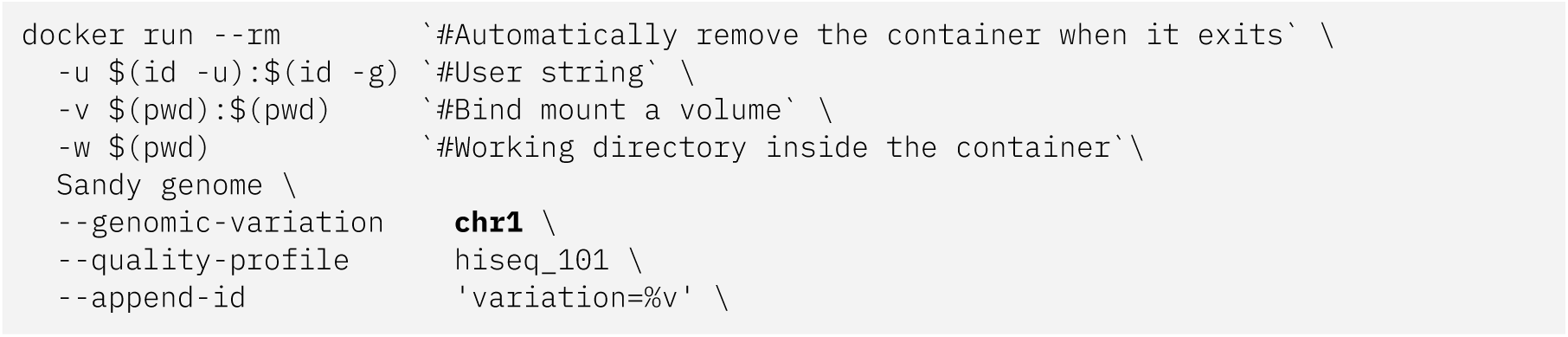

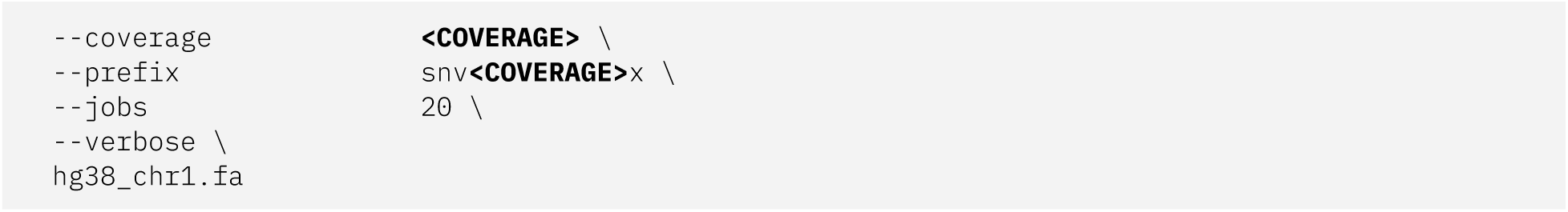

We added variation tags for the reads simulated using the variant (`append_id`). Then, we mapped BWA using only hg38 chromosome 1 in the BWA MEM:

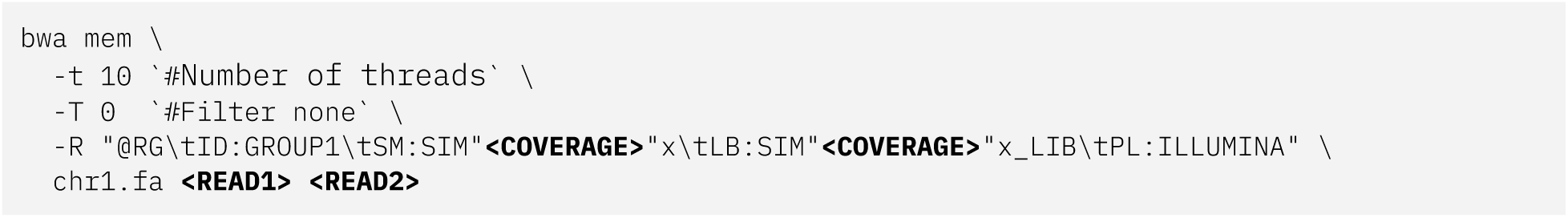

Next, low-alignment score reads (Q < 20) and potential PCR duplicates (using Picard’s Mark Duplicate) were removed. We rescored our alignment using GATK’s Base Recalibrator (https://gatk.broadinstitute.org/) and the GATK resource bundle `(knownSites dbsnp_146. hg38.vcf` and `knownSites Mills_and_1000G_gold_standard.indels.hg38.vcf`). Variant calling was performed with GATK’s Haplotype Caller using the following parameters:

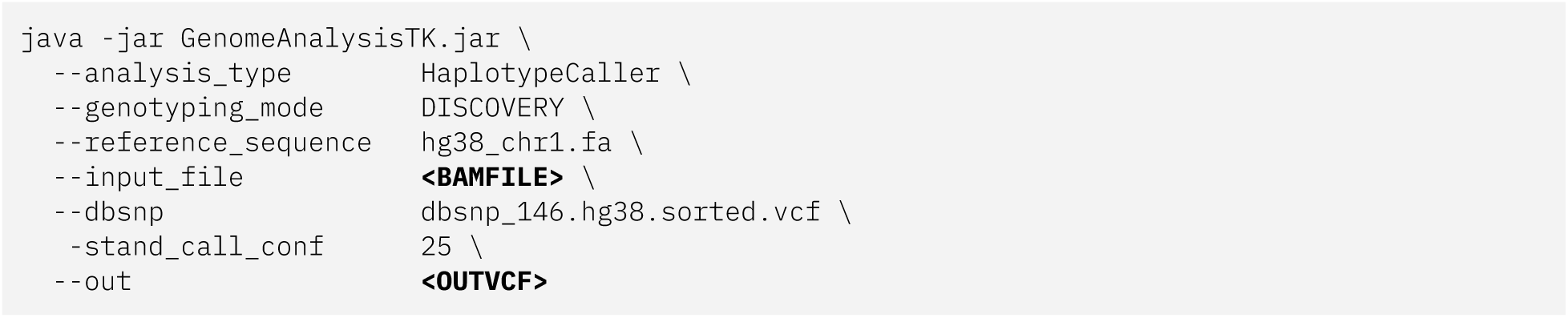

Finally we split the biallelic sites and filtered data using GATK’s recommend Hard Filtering (QD < 2.0 || FS > 60.0 || MQ < 40.0 || MQRankSum < −12.5 || ReadPosRankSum < −8.0 for SNVs, and QD < 2.0 || FS > 200.0 || ReadPosRankSum < −20.0 for indels).

### 4.6. Figures

Figures were constructed using R (https://www.r-project.org/), Circos (Krzywinski et al. 2009), and Inkscape (https://inkscape.org/).

## Availability of data and materials

All data used in this study are publicly available (access IDs: SRR5805510, SRR3185389, SRR3355336, SRR6876696, SRR7089434, SRR6131534, SRR544541, PRJEB23027, SRP012400, SRP000547, SRP018096, SRP000032, and ERP001229) at the European Nucleotide Archive (https://www.ebi.ac.uk/ena) or Short Read Archive (https://www.ncbi.nlm.nih.gov/sra).

## Acknowledgments

We thank Anamaria A. Camargo and Cibele Masotti for their helpful comments. We also thank the members of the Galante laboratory for their comments and testing of Sandy’s pipeline. This work was partially supported by Fundação de Amparo à Pesquisa do Estado de São Paulo (2018/15579-8), Instituto Serrapilheira, and Conselho Nacional de Desenvolvimento Científico e Tecnológico (CNPq to PAFG). GDAG (2017/19541-2), FRCS (2017/18246-7), FORR (2015/25020-0), HBC (2018/13613-4), and RLVM (2020/02413-4) were supported by fellowships from the Fundação de Amparo à Pesquisa do Estado de São Paulo (FAPESP). TAM is supported by a fellowship from Coordenação de Aperfeiçoamento de Pessoal de Nível Superior (CAPES). JLB and RASB received fellowships from CNPq.

## Author Contributions

TAM contributed to the study design, development, implementation, data analysis, online documentation, figures, and manuscript preparation. GDAG contributed to the analyses of Sandy’s performance, gene expression, SNVs calling, fusion detection, figures, and manuscript preparation. HBC contributed to the analysis of Sandy’s performance, gene expression, figures, and manuscript preparation. FRCS contributed to the analysis of Sandy’s performance, gene expression, figures, and manuscript preparation. RASB contributed to the SNV detection analysis, manuscript preparation, and figures. JLB contributed to online documentation and manuscript preparation. FORR contributed to study design, figures, and manuscript preparation. PAFG coordinated the study and prepared the manuscript with input from other authors. All the authors have read and approved the final version of the manuscript.

