## Supplemental Figures and Tables for "Sandy: A user-friendly and versatile NGS simulator to facilitate sequencing assay design and optimization"

Thiago L. A. Miller[1,2], Gabriela Guardia[1], Helena B. Conceição[1,3], Rafael L. Mercuri[1,3], Felipe R. C. Santos[1,3], Rodrigo Barreiro[1,2], José Leonel Buzzo[1,2], Fernanda O. Rego[1] & Pedro A. F. Galante[1]
1 - Hospital Sirio Libanes, São Paulo, Brazil.
2 - Instituto de Química, Dept de Bioquímica, Universidade de São Paulo, Brazil
3 - Interunidades em Bioinformática, Universidade de São Paulo, Brazil

### Supplementary Data

```
$ for tissue in stomach testis pancreas brain_hippocampus; do
  docker run \
  --rm \
  -v "$PWD/sandy:/mnt" \
  -v "gencode.v36.transcripts.fa.gz:/data/gencode.fa.gz:ro" \
  -u "$(id -u):$(id -g)" \
        galantelab/sandy:latest
                transcriptome \
                --verbose \
                --jobs=5 \
                --seed=1717 \
                --output-dir=/mnt/$tissue/100 \
                --number-of-reads=100000000 \
                --expression-matrix=$tissue \
                --sequencing-type=paired-end \
                --quality-profile=hiseq_76 \
                --fragment-mean=237 \
                --fragment-stdd=82 \
                /data/gencode.fa.gz \
        '2>' $PWD/log/$tissue_100.log \
  done
```

Figure S1. Command line used to simulate expression in the stomach, testis, pancreas, and hippocampus (dataset used to build Figure 3C).

```
$ #Run RNA-seq simulations for Illumina HiSeq  in server ##################
       for reads in 10 20 30 40 50 60 70 80 90 100 110 120 130 140 150; do
       { time docker run --rm -u $(id -u):$(id -g) -v $(pwd -P):/mnt -w /mnt \
             galantelab/sandy:dev  transcriptome  -v  -q  hiseq_101  -f  liver  -n  ${reads}000000
gencode.v40.transcripts.fa.gz \
              --jobs=20 --seed 123; } 2> transcriptome${reads}_time.out
       done

#Run RNA-seq simulations for Illumina HiSeq in laptop ##################
       for reads in 10 20 30 40 50 60 70 80 90 100; do
       { time docker run --rm -u $(id -u):$(id -g) -v $(pwd -P):/mnt -w /mnt \
             galantelab/sandy:dev  transcriptome  -v  -q  hiseq_101  -f  liver  -n  ${reads}000000
gencode.v40.transcripts.fa.gz \
              --jobs=4 --seed 123; } 2> transcriptome${reads}_time.out
       done
```

Figure S2. Command line used to simulate RNA-Seq using a *server* and *laptop* (dataset used to build Figure 5A and 5F).

```
$ #Run WGS simulations for Illumina HiSeq in server ###################
        for cov in 1 2 4 8 16 32 64 128; do
                { time docker run --rm -u $(id -u):$(id -g) -v $(pwd -P):/mnt -w /mnt \
                    galantelab/sandy:dev genome -v -q hiseq_101 -c $cov hg38.fa \
                        --jobs=20 --seed 123; } 2> hiseq${cov}_time.out
        done

#Run WGS simulations for Illumina HiSeq in laptop ###################
        for cov in 1 2 4 8 16; do
                { time docker run --rm -u $(id -u):$(id -g) -v $(pwd -P):/mnt -w /mnt \
                    galantelab/sandy:dev genome -v -q hiseq_101 -c $cov hg38.fa \
                        --jobs=4 --seed 123; } 2> hiseq${cov}_time.out
        done

#Run WGS simulations for Oxford Nanopore in server ###################
        for cov in 1 2 4 8 16 32 64 128; do
                { time docker run --rm -u $(id -u):$(id -g) -v $(pwd -P):/mnt -w /mnt \
                    galantelab/sandy:dev genome -v -q ont -c $cov hg38.fa \
                        --jobs=20 --seed 123; } 2> oxford${cov}_time.out
        done

#Run WGS simulations for Oxford Nanopore in laptop ###################
        for cov in 1 2 4 8 16; do
                { time docker run --rm -u $(id -u):$(id -g) -v $(pwd -P):/mnt -w /mnt \
                    galantelab/sandy:dev genome -v -q ont -c $cov hg38.fa \
                        --jobs=4 --seed 123; } 2> oxford${cov}_time.out
        done

#Run WGS simulations for PacBio in server ###################
        for cov in 1 2 4 8 16 32 64 128; do
                { time docker run --rm -u $(id -u):$(id -g) -v $(pwd -P):/mnt -w /mnt \
                    galantelab/sandy:dev genome -v -q pacbio -c $cov hg38.fa \
                        --jobs=20 --seed 123; } 2> pacbio${cov}_time.out
        done

#Run WGS simulations for PacBio in laptop ###################
        for cov in 1 2 4 8 16; do
                { time docker run --rm -u $(id -u):$(id -g) -v $(pwd -P):/mnt -w /mnt \
                    galantelab/sandy:dev genome -v -q pacbio -c $cov hg38.fa \
                        --jobs=4 --seed 123; } 2> pacbio${cov}_time.out
```

```
        done

#Run genome simulations for Illumina HiSeq with variations in server ###################
        for cov in 1 2 4 8 16 32 64 128; do
        #Run genome simulations for Illumina HiSeq with SNVs/INDELs ###################
        { time docker run --rm -u $(id -u):$(id -g) -v $(pwd -P):/mnt -w /mnt \
            galantelab/sandy:dev genome -v -q hiseq_101 -c $cov -A NA12878 hg38.fa \
                --jobs=20 --seed 123; } 2> hiseq${cov}SNVs_time.out
        echo "Hiseq SNVs $cov ALL DONE"

        #Run genome simulations for Illumina HiSeq with FUSIONS #######################
        { time docker run --rm -u $(id -u):$(id -g) -v $(pwd -P):/mnt -w /mnt \
            galantelab/sandy:dev genome -v -q hiseq_101 -c $cov -A fusion hg38.fa \
                --jobs=20 --seed 123; } 2> hiseq${cov}fusion_time.out
        echo "Hiseq Fusion $cov ALL DONE"

        #Run genome simulations for Illumina HiSeq with SNVs/INDELs & FUSIONS ########
        { time docker run --rm -u $(id -u):$(id -g) -v $(pwd -P):/mnt -w /mnt \
            galantelab/sandy:dev genome -v -q hiseq_101 -c $cov -A NA12878 -A fusion hg38.fa \
                --jobs=20 --seed 123; } 2> hiseq${cov}allSV_time.out
        echo "Hiseq Fusion & SNVs $cov ALL DONE"
        done
```

Figure S3. Command line used to simulate Whole Genome Sequencing using a *server* and *laptop* (dataset used to build Figure 5B-F).

```
$  for tissue in bladder brain_cortex brain_hippocampus liver pancreas stomach; do
       for coverage in 10 30 50 100 150; do
         docker run \
         --rm \
         -v "$PWD/sandy:/mnt" \
         -v "gencode.v36.transcripts.fa.gz:/data/gencode.fa.gz:ro" \
         -u "$(id -u):$(id -g)" \
               galantelab/sandy:latest
                       transcriptome \
                       --verbose \
                       --jobs=5 \
                       --seed=1717 \
                       --output-dir=/mnt/$tissue/100 \
                       --number-of-reads=${coverage}000000 \
                       --expression-matrix=$tissue \
                       --sequencing-type=paired-end \
                       --quality-profile=hiseq_76 \
                       --fragment-mean=237 \
                       --fragment-stdd=82 \
                       /data/gencode.fa.gz \
               '2>' $PWD/log/$tissue_${coverage}.log \
       done
  done
```

Figure S4. Command line used to simulate RNA-seq based on the expression pattern of Bladder, Brain Cortex, Brain Hippocampus, Liver, Pancreas and Stomach for coverages of 10 30 50 100 150 (dataset used to build Figure 6A-H).

```
$ for coverage in 4 8 16 32 64 128; do
  docker run --rm -u $(id -u):$(id -g) -v $(pwd -P):/mnt -w /mnt galantelab/sandy genome \
      --verbose \
      --prefix=SNVs_${coverage} \
      --output-format=fastq.gz \
      --jobs=20 \
      --append-id='variation=%v' \
      --quality-profile=hiseq_101 \
      --sequencing-type=paired-end \
      --fragment-mean=300 \
```

```
      --fragment-stdd=50 \
      --coverage=${coverage} \
      --genomic-variation=NA12878_hg38_chr1 \
      hg38.chr1.fa.gz;
  done
```

Figure S5. Command line used to simulate human genome chromosome 1 with SNVs and indels from the individual NA12878 at different coverages (4X, 8X, 16X, 32X, 64X, and 128X) (dataset used to build Figure 6A-B).

```
$ for coverage in 4 8 16; do
  docker run --rm -u $(id -u):$(id -g) -v $(pwd -P):/mnt -w /mnt galantelab/sandy genome \
      --verbose \
      --prefix=fusions_${coverage} \
      --output-format=fastq.gz \
      --jobs=20 \
      --append-id='variation=%v' \
      --quality-profile=hiseq_101 \
      --sequencing-type=paired-end \
      --fragment-mean=300 \
      --fragment-stdd=50 \
      --coverage=${coverage} \
      --genomic-variation-regex=fusion \
      hg38.fa.gz;
  done
```

Figure S6. Command line used to simulate human genome with 10 homozygous gene fusions at different coverages (4X, 8X and 16X)  (dataset used to build Figure 6C).

#### Supplementary tables

Table S1. Genomic position of 10 cases of homozygous gene fusions reported by COSMIC.

| | Gene 1 | | | | | Gene 2 | | | | |
|---|---|---|---|---|---|---|---|---|---|---|
| gene.fusion | strand | chr | start | fusion.point | end | strand | chr | start | fusion.point | end |
| EML4-ALK | + | chr2 | 42169350 | 42283588 | 42332548 | - | chr2 | 29192774 | 29411411 | 29921566 |
| KIAA1549-BRAF | - | chr7 | 138831381 | 138936336 | 138981318 | - | chr7 | 140719327 | 140781121 | 140924928 |
| NCOA4-RET | - | chr10 | 46005088 | 46023025 | 46030714 | + | chr10 | 43077027 | 43093023 | 43130351 |
| CCDC6-RET | - | chr10 | 59788763 | 59871287 | 59906656 | + | chr10 | 43077027 | 43093023 | 43130351 |
| TMPRSS2-ERG | - | chr21 | 41464551 | 41511146 | 41531116 | - | chr21 | 38380027 | 38464552 | 38661780 |
| NPM1-ALK | + | chr5 | 171387116 | 171403930 | 171411137 | - | chr2 | 29192774 | 29411411 | 29921566 |
| KMT2A-AFF1 | + | chr11 | 118436490 | 118499728 | 118526832 | + | chr4 | 86935002 | 86996817 | 87141054 |
| BCR-ABL1 | + | chr22 | 23179704 | 23276536 | 23318037 | + | chr9 | 130713946 | 130766064 | 130887675 |
| EWSR1-ERG | + | chr22 | 29268009 | 29290769 | 29300525 | - | chr21 | 38380027 | 38464541 | 38661780 |
| EWSR1-FLI1 | + | chr22 | 29268009 | 29290769 | 29300525 | + | chr11 | 128686535 | 128724554 | 128813267 |

Table S2. Comparison of Simulation Coverage for Fusion Detection

| | chr1 | chr1.pos | chr2 | chr2.pos | type | filter | precision | PE | MAPQ | SR | Detected at 4x | Detected at 8x | Detected at 16x |
|---|---|---|---|---|---|---|---|---|---|---|---|---|---|
| EML4-ALK | chr2 | 29921566 | chr2 | 42332600 | deletion | PASS | IMPRECISE | 3 | 60 | 0 | X | X | X |
| KIAA1549-BRAF | chr7 | 138936336 | chr7 | 140780958 | deletion | PASS | PRECISE | 4 | 60 | 4 | X | X | X |
| NCOA4-RET | chr10 | 43130351 | chr10 | 46030756 | deletion | PASS | IMPRECISE | 3 | 60 | 0 | X | X | X |
| CCDC6-RET | chr10 | 43130336 | chr10 | 59906656 | deletion | PASS | IMPRECISE | 4 | 60 | 0 | X | X | X |
| TMPRSS2-ERG | chr21 | 38661779 | chr21 | 41531116 | deletion | PASS | PRECISE | 4 | 60 | 8 | | X | X |
| BCR-ABL1 | chr9 | 130766322 | chr22 | 23276351 | translocation | PASS | IMPRECISE | 5 | 60 | 0 | | X | X |
| EWSR1-ERG | chr22 | 29300525 | chr21 | 38661779 | translocation | PASS | PRECISE | 6 | 60 | 8 | | X | X |
| NPM1-ALK | chr5 | 171411415 | chr2 | 29921374 | translocation | PASS | IMPRECISE | 5 | 60 | 0 | | | X |
| KMT2A-AFF1 | chr4 | 86996817 | chr11 | 118499727 | translocation | PASS | PRECISE | 9 | 60 | 6 | | | X |
| EWSR1-FLI1 | chr22 | 29290769 | chr11 | 128724555 | translocation | PASS | PRECISE | 8 | 60 | 8 | | | X |

PASS: All filters passed
PRECISE: has split-read support (SR>0)
PE: Paired-end support of the structural variant
MAPQ: Median mapping quality of paired-ends
SR: Split-read support
